# A *pcyt-1* Allelic Series Reveals In Vivo Consequences of Reduced Phosphatidylcholine Synthesis in *C. elegans*

**DOI:** 10.64898/2026.04.22.720214

**Authors:** August Qvist, Delaney Kaper, Marcus Henricsson, Albin Stjernman, Jan Borén, Marc Pilon

**Affiliations:** Department of Chemistry and Molecular Biology, University of Gothenburg, 405 30 Gothenburg, Sweden and; Department of Molecular and Clinical Medicine/Wallenberg Laboratory, Institute of Medicine, University of Gothenburg, 405 30 Gothenburg, Sweden

## Abstract

Phosphatidylcholine (PC) is the most abundant phospholipid in eukaryotic membranes and is synthesized in part via the rate-limiting enzyme PCYT1A. In humans, hypomorphic PCYT1A variants cause diverse disorders, including retinal dystrophy, lipodystrophy with fatty liver, and spondylometaphyseal dysplasia. To define how graded reductions in PC synthesis affect organismal physiology, we generated and characterized a series of mutant alleles in the *Caenorhabditis elegans* homolog *pcyt-1*, including variants corresponding to disease-causing human mutations, as well as an auxin-inducible degradation (AID) allele.

We identify a clear allelic hierarchy. The V146M variant is embryonic lethal, whereas A97T is largely benign. P154A is temperature-sensitive, and C211Y causes growth delay, reduced brood size, sterility, and lengthened lifespan at standard temperature. Phenotypes of C211Y are rescued by choline, CDP-choline, or phosphatidylcholine supplementation, supporting reduced enzymatic function. Lipidomic profiling reveals that decreased PC synthesis consistently increases long-chain polyunsaturated fatty acids (LCPUFAs) in both PCs and PEs at the expense of shorter saturated species, without markedly altering the PC/PE ratio at 20°C. At elevated temperature, the P154A variant exhibits protein instability and a decreased PC/PE ratio.

Despite significant lipid remodeling, canonical ER, mitochondrial, and metabolic stress GFP-based reporters are not activated; only the oxidative stress response is elevated, consistent with increased peroxidation-prone LCPUFAs in the *pcyt-1* mutant. Acute auxin-induced degradation of PCYT-1 in larvae causes developmental arrest, while acute PCYT-1 degradation in adults disrupts oogenesis, demonstrating a continuous requirement for PC synthesis. Together, these findings establish a functional *pcyt-1* allelic series and show that limiting PC synthesis drives compensatory remodeling toward LCPUFA-enriched membranes while rendering the germline particularly vulnerable.

## INTRODUCTION

Homodimeric PCYT1A is the rate-limiting enzyme in the synthesis of phosphatidylcholine (PC), the most abundant phospholipid in most eukaryotic cellular membranes **(Fig. 1A)**. Low PC levels lead to packing defects in the inner nuclear membranes that allow recruitment and activation of PCYT1A, thus leading to increased PC synthesis until the membrane defects are corrected **(Fig. 1B)** (Haider *et al*. 2018). PCYT1A is essential in mammals, and so is its homolog *pcyt-1* in the nematode *C. elegans*. However, several alleles of PCYT1A are viable in mammals and cause a variety of conditions; reviewed in (Cornell *et al*. 2019) (**Fig. 1C)**. Some missense mutations (e.g. A93T) cause retinal dystrophies, specifically Leber Congenital Amaurosis, abbreviated as LCA (Testa *et al*. 2017). Other missense variants (e.g. V142M) cause lipodystrophy and fatty liver disease in human, abbreviated as CLD-FL (Payne *et al*. 2014). Finally, some missense mutations (e.g. P150A) cause spondylometaphyseal dysplasia (a form of dwarfism) in human (Hoover-Fong *et al*. 2014; Wong 2014; Yamamoto *et al*. 2014) and dogs (Ludwig-Peisker *et al*. 2022), accompanied at least in human with cone-rod dystrophy; these defects are collectively designated SMD-CRD. In the budding yeast *S. cerevisiae,* PC deficiency in a choline auxotroph can be compensated by mutations that cause shortened FA chains, suggesting that a primary function of PC in eukaryotic cells is to promote membrane fluidity (Boumann *et al*. 2006; Bao *et al*. 2021).

**Fig. 1.**
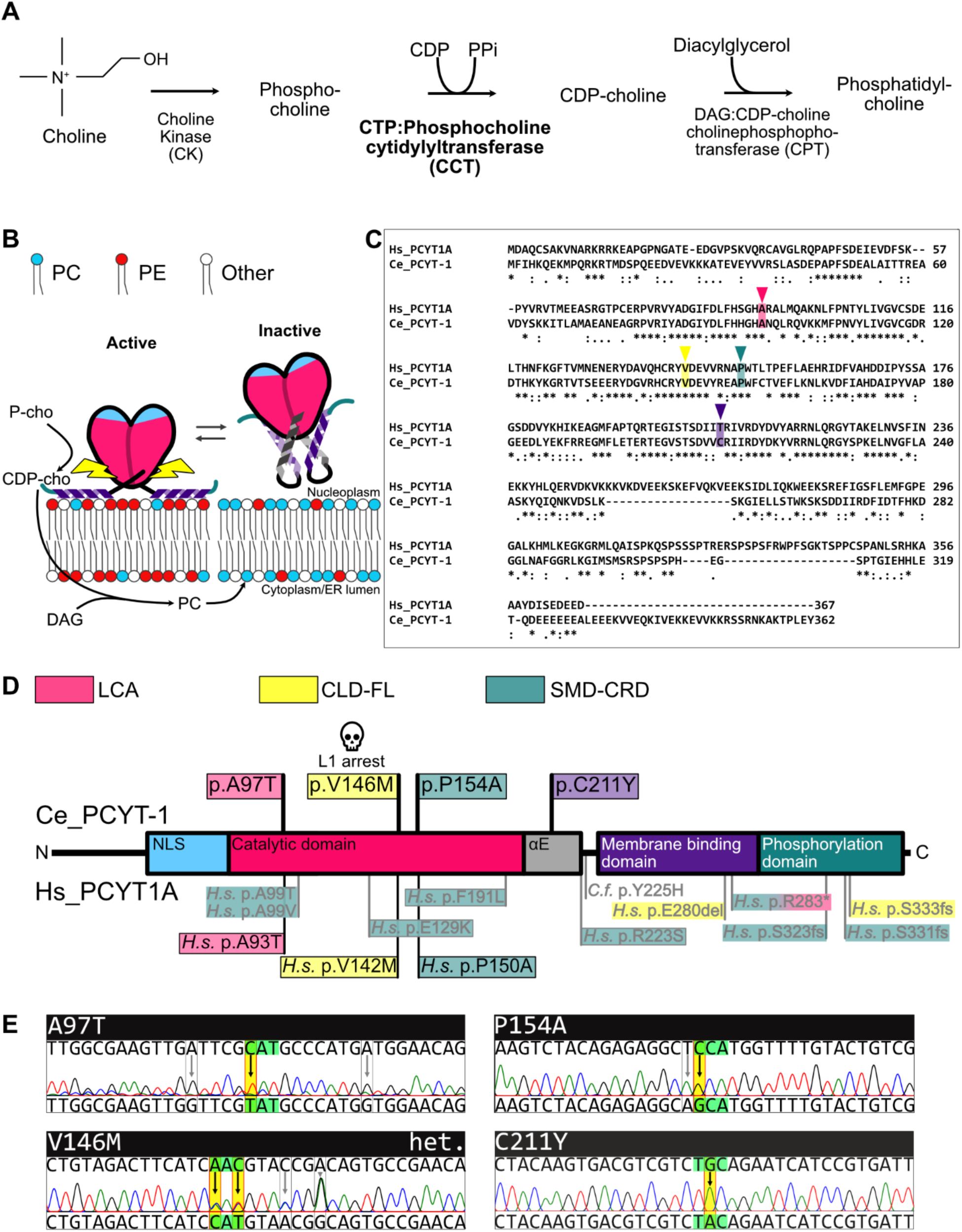
Hs PCYT1A and *Ce* PCYT-1 function and mutant alleles. **A)** Phosphatidylcholine branch of the Kennedy pathway. **B)** Schematic of the regulation of CCT inside the nucleus. **C)** Amino acid alignment of human PCYT1A and *C. elegans* PCYT-1.* = Fully conserved residues, : = Conservation between groups of strongly similar properties,. = Conservation between groups of weakly similar properties. **D)** Protein schematic PCYT1A/PCYT-1 highlighting missense mutations in *C. elegans* PCYT-1 and disease-causing mutations in human and dog PCYT1A. NLS=nuclear localization signal, αE is the alpha helix forming the hinge region between the catalytic and the membrane binding domains. **E)** Nucleotide changes and DNA sequence trace of the three CRISPR/Cas9 generated disease homologous mutants A97T, P154A and V146M, and the C211Y mutant in *C. elegans*. Note that V146M is from a heterologous sample.

The nematode *C. elegans* is well suited for the genetic analysis of lipid metabolism not least because it is capable of synthesizing all of its essential fatty acids, including long-chain polyunsaturated fatty acids (LCPUFAs) via thoroughly described pathways (Watts and Ristow 2017), and because of the ease with which forward genetic studies via random mutagenesis (Watts and Browse 2002; Svensk *et al*. 2013; Devkota *et al*. 2021a; Kaper *et al*. 2025) or reverse genetic studies via CRISPR/Cas9 genome editing (Friedland *et al*. 2013; Dickinson *et al*. 2015; Paix *et al*. 2017; Dokshin *et al*. 2018) can be performed.

How PC homeostasis impacts *C. elegans* has also been studied. PC deficiency increases ferroptosis susceptibility in the germline, leading to sterility (Zhu *et al*. 2023). Also, PC supplements prevent starvation-induced fertility defects (Jordan *et al*. 2023) and extend lifespan via DAF-16 while reducing amyloid-beta-induced toxicity (Kim *et al*. 2019). Diet can also impact PC levels in *C. elegans*; for example, cultivating the dietary *E. coli* bacteria in the presence of glucose results in decreased PC levels accompanied by increased saturated fatty acids (SFAs) and cyclopropane fatty acids in the worms (Vieira *et al*. 2022; Wang *et al*. 2025), as well as reduced fecundity. Unfortunately, few *pcyt-1* alleles have been characterized until now. One, the *pcyt-1(et9)* allele, was isolated as a *paqr-*2 suppressor, suggesting that it may act by promoting membrane fluidity (Svensk *et al*. 2013), though the precise mechanism remains ill-defined. *pcyt-1(et9)* also causes increased ferroptosis in the germline as a single mutant (Zhu *et al*. 2023), while suppressing the embryonic lethality of *seip-1(tm4221)*, a seipin null allele (Zhu *et al*. 2022); both effects may be due to *pcyt-1(et9)* causing increased PUFA levels in PCs though the reason for this also remains ill-defined.

To better understand how varying degrees of PC deficiency can impact an organism, we created a small collection of *pcyt-1* alleles in *C. elegans*, including a variant of which the degradation of PCYT-1 is auxin-inducible, evaluated their phenotypes and determined the fatty acid composition of their phosphatidylcholines and phosphatidylethanolamines. In this way we define a useful *pcyt-1* allelic series and show that disrupting PC synthesis increases the abundance of PUFAs in both PCs and PEs.

## RESULTS

Using CRISPR/Cas9, we created mutations in the *C. elegans pcyt-1* gene that corresponds to missense mutations causing three distinct disorders in humans and that have previously been biochemically characterized in overexpression assays (Cornell *et al*. 2019): A97T, V146M and P154A correspond to the human A93T, V142M and P150A that cause LCA, CLD-FL and SMD-CRD, respectively (**Fig. 1C-E**). Additionally, we also studied the *pcyt-1(et9)* single mutant that causes a C211Y missense within or near the hinge regulatory region (Cornell *et al*. 2019) (**Fig. 1C-E**). For clarity, we will designate the *pcyt-1* alleles by their missense mutations (e.g., *pcyt-1(C211Y)* or simply C211Y) throughout this article.

### A hierarchy of *pcyt-1* variants

Of the studied variants, the V146M is the most severe since worms homozygous for it arrested as eggs. The other variants were viable and displayed a hierarchy of effects on several traits. At 15°C and 20°C, only the C211Y variant caused a significant growth defect when the length of worms was measured 144 or 72 hours respectively after placing synchronized L1 larvae on culture plates (**Fig. 2A-B**). At 25°C, both the P154A and C211Y variants caused a growth defect (**Fig. 2C**). This was the first indication that the P154A is a temperature-sensitive variant. At 15°C, none of the *pcyt-1* allele exhibit any sterility phenotype (**Fig. 2D**). At 20°C, only C211Y cause sterility in ∼40% of the worms (**Fig. 2E**). At 25°C, both the P154A and C211Y alleles show a very high penetrance (>90%) of sterility (**Fig. 2F**). Interestingly, the sterility phenotype in *pcyt-1* mutants is binary: either a worm seems to have normal gonads and number of eggs in its uterus, or it has gonad arms that are uniformly devoid of oocytes, as shown in **Fig. 2G**. Lifespan was also evaluated: C211Y caused an extension of lifespan at 20°C (**Fig. 2H**), and both P154A and C211Y extended lifespan at 25°C (**Fig. 2I**). Brood size was determined at 20°C; of the alleles studied, only C211Y caused a significant reduction in the number of viable progeny produced (**Fig. 2J**). We conclude that C211Y is a relatively severe allele at 20°C while the A97T and P154A variants are mostly harmless at that temperature. Interestingly, P154A is a temperature-sensitive allele: at 25°C, both the C211Y and P154A are equally severe variants, while again A97T appears harmless.

**Fig. 2.**
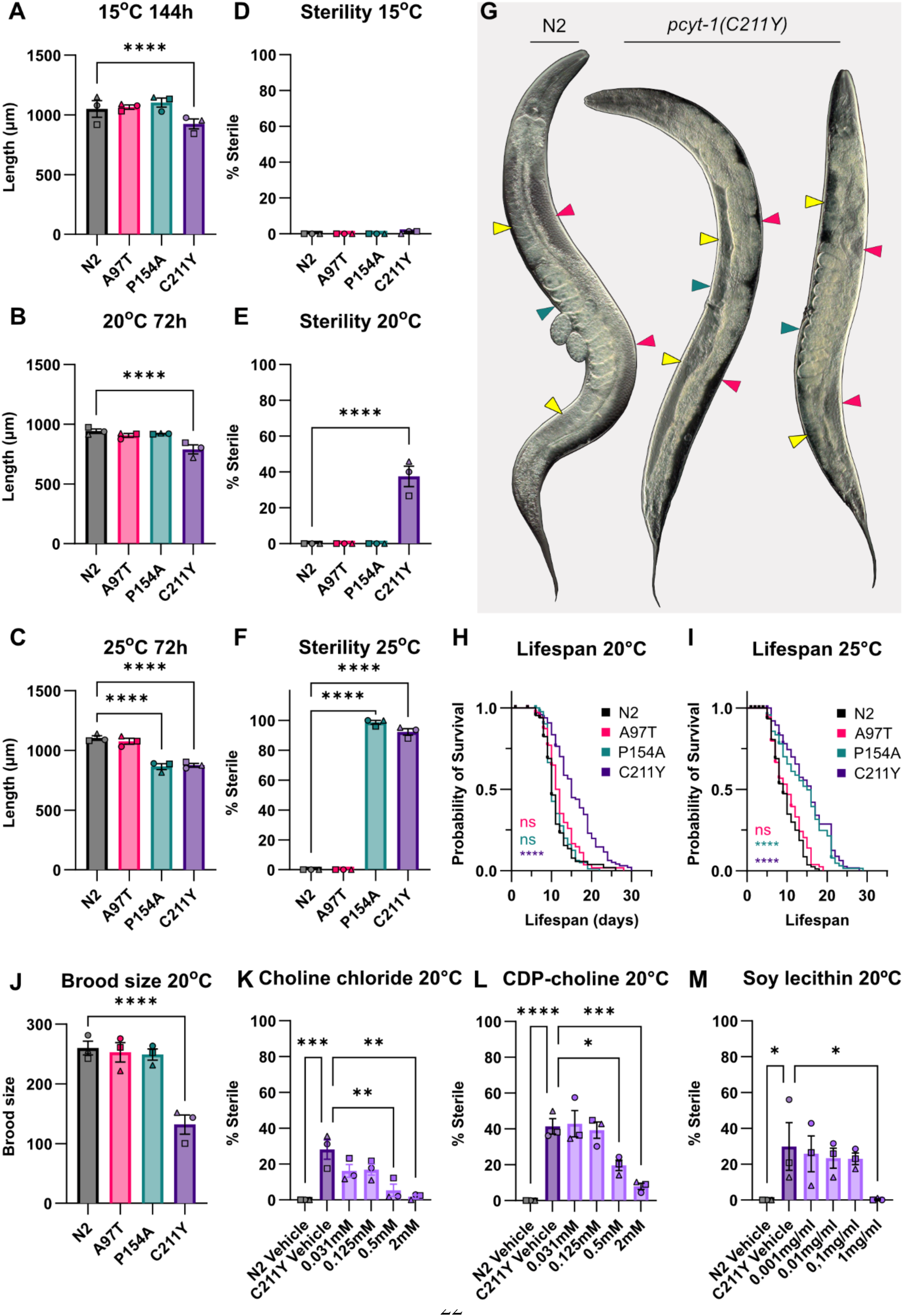
Growth, sterility and lifespan of pcyt-1 mutant alleles at different temperatures. A-. **C)** Length measurements, and (**D-F)** fraction of sterile worms, at 15°C after 144h and at 20°C and 25°C after 72h. **G)** Green arrows point to the uterus of the worm, yellow arrows point to the proximal side of the gonad where oocytes are located, and pink arrows point to the distal side of the gonad where the germ line is located. For the sterile worm, the arrows point to these locations where there would normally be eggs, oocytes, and germ cells. Note how N2 has clearly visible and a normal amount of nicely aligned oocytes and normal-looking germline, while the fertile C211Y has fewer visible oocytes and less distinguished germline. The sterile C211Y has a completely empty gonad, with no visible oocytes or germline. **H-I)** Lifespan at 20°C and 25°C. **J)** Total brood size at 20°C. **K-M)** Sterility of C211Y worms after 72h when treated with Choline chloride (K), CDP-choline (L), or Soy lecithin (M). A-C, J) Two-way mixed-effects ANOVA (REML) using LS means (GraphPad Prism), with biological replicate treated as a random effect. Tukey’s multiple comparisons test was used for post-hoc analysis, comparing each strain to N2. A-C) n = 3 biological replicates (19–25 worms per replicate). J) n = 3 biological replicates (5 worms per replicate). D-F, K-M) Ordinary one-way ANOVA was used, with Dunnett’s multiple comparisons test for post-hoc analysis comparing each strain to N2 (D-F), or each tested concentration to C211Y Vehicle (K-M). n=3 biological replicates with ≥30 worms per replicate. * == p<0,5; ** == p<0,05; *** == p<0,005; **** == p<0,0005.

Importantly, the phenotypes of the C211Y allele can be suppressed by the addition of choline (a substrate of the PCYT-1 reaction), CDP-choline (the product of the PCYT-1 reaction) or soy lecithin (rich in PCs), as exemplified by the suppression of sterility at 20°C (**Fig. 2K-M**). We conclude that C211Y is likely a reduction-of-function allele, i.e., that the mutated protein has reduced function, and that the essential output is the synthesis of PCs.

The C211Y variant was isolated in a forward genetic screen for suppressors of the cold intolerance and tail end defect found in the *paqr-2* mutant (Svensk *et al*. 2013). *paqr-2*, a homolog of the human AdipoR1 and AdipoR2, is essential for membrane fluidity homeostasis especially in response to low temperature or SFA-rich diets (Svensk *et al*. 2013; Svensk *et al*. 2016; Devkota *et al*. 2017; Devkota *et al*. 2021a; Devkota *et al*. 2021b; Ruiz *et al*. 2022; Pilon and Ruiz 2023; Ruiz *et al*. 2023). Here, we found that only C211Y suppressed the *paqr-2* tail end defect at 20°C, while both P154A and C211Y suppressed it at 25°C, confirming again that P154A is temperature-sensitive variant (**Fig. 3A-C**). The A97T and P154A variants can significantly suppress the cold intolerance defect of the *paqr-2* mutant though the effect size is not as large as the C211Y variant (**Fig. 3D**). Conversely, *paqr-2* did not suppress the sterility of the PCYT-1 P154A and C211Y variants (**Fig. 3E**), suggesting that the suppression is not mutual.

**Fig. 3.**
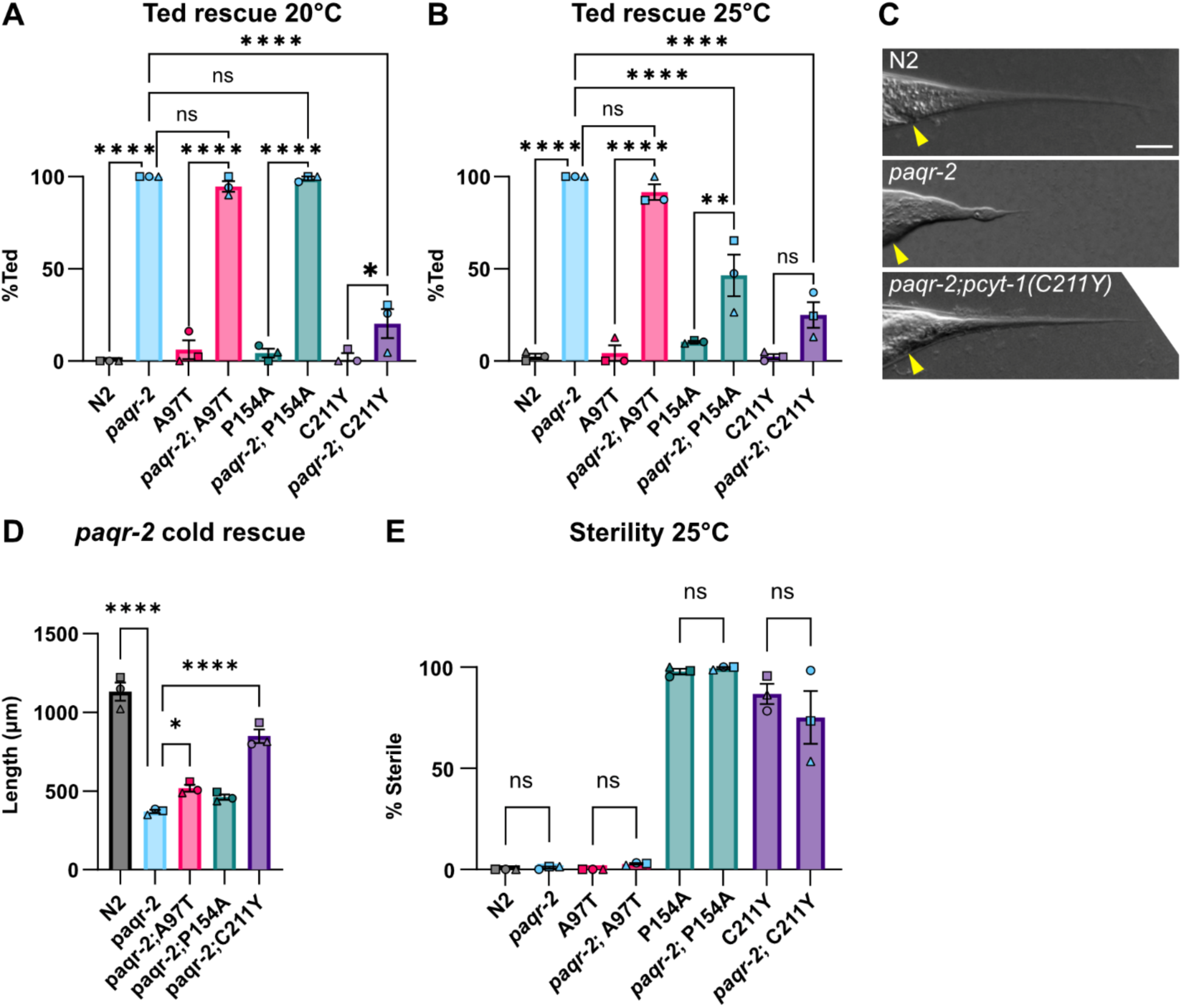
Genetic interactions between the *paqr-2(tm3410)* null and different *pcyt-1* mutant alleles. **A-B)** Fraction of day 1 adults with a Tail End Defective in the hermaphrodite (Ted) phenotype. **C)** Representative images of the tail tip in N2 *paqr-*2 and the *paqr-2;pcyt-1(C211Y)* double mutant where *paqr-*2 presents a severe Ted phenotype. Scalebar = 25 micrometer, yellow arrow points to anus. **D)** Length measurements of *paqr-2* and *pcyt-1* double mutants after 144h at 15°C. **E)** Fraction of sterile worms after 72h at 25°C. A-B, E) Ordinary one-way ANOVA was used, with Dunnett’s multiple comparisons test for post-hoc analysis comparing each strain (D-F) to N2, or (K-M) each tested concentration to C211Y Vehicle. n=3 biological replicates with ≥30 worms per replicate. D) Two-way mixed-effects ANOVA (REML) using LS means (GraphPad Prism), with biological replicate treated as a random effect. Tukey’s multiple comparisons test was used for post-hoc analysis, comparing to each strain to *paqr-2*. n = 3 biological replicates (19–25 worms per replicate). * == p<0,5; ** == p<0,05; *** == p<0,005; **** == p<0,0005

### Stress responses are subdued in the *pcyt-1(C211Y)* mutant

Among the variable *pcyt-1* variants studied, the C211Y variant has the strongest phenotypic effect at 20°C, the standard laboratory condition. One may therefore expect that stress response pathways related to membrane-bound organelles or metabolism would be constitutively activated in this mutant. However, we found that only the oxidative stress response pathway, assessed using a *gst-4::GFP* reporter (Link and Johnson 2002)(**Fig. 4A-B**), is upregulated while the metabolic stress response *(daf-16::GFP;* (Libina et al. 2003))(**Fig. 4C-D**), endoplasmic reticulum unfolded protein response (*hsp-4::GFP;* (Calfon *et al*. 2002))(**Fig. 4E-F**), and mitochondrial unfolded protein response (*hsp-60::GFP;* (Yoneda et al. 2004)) (**Fig. 4G-H**) pathways are not. This suggests reduced PC synthesis is generally well tolerated but leads to increased oxidative stress.

**Fig. 4.**
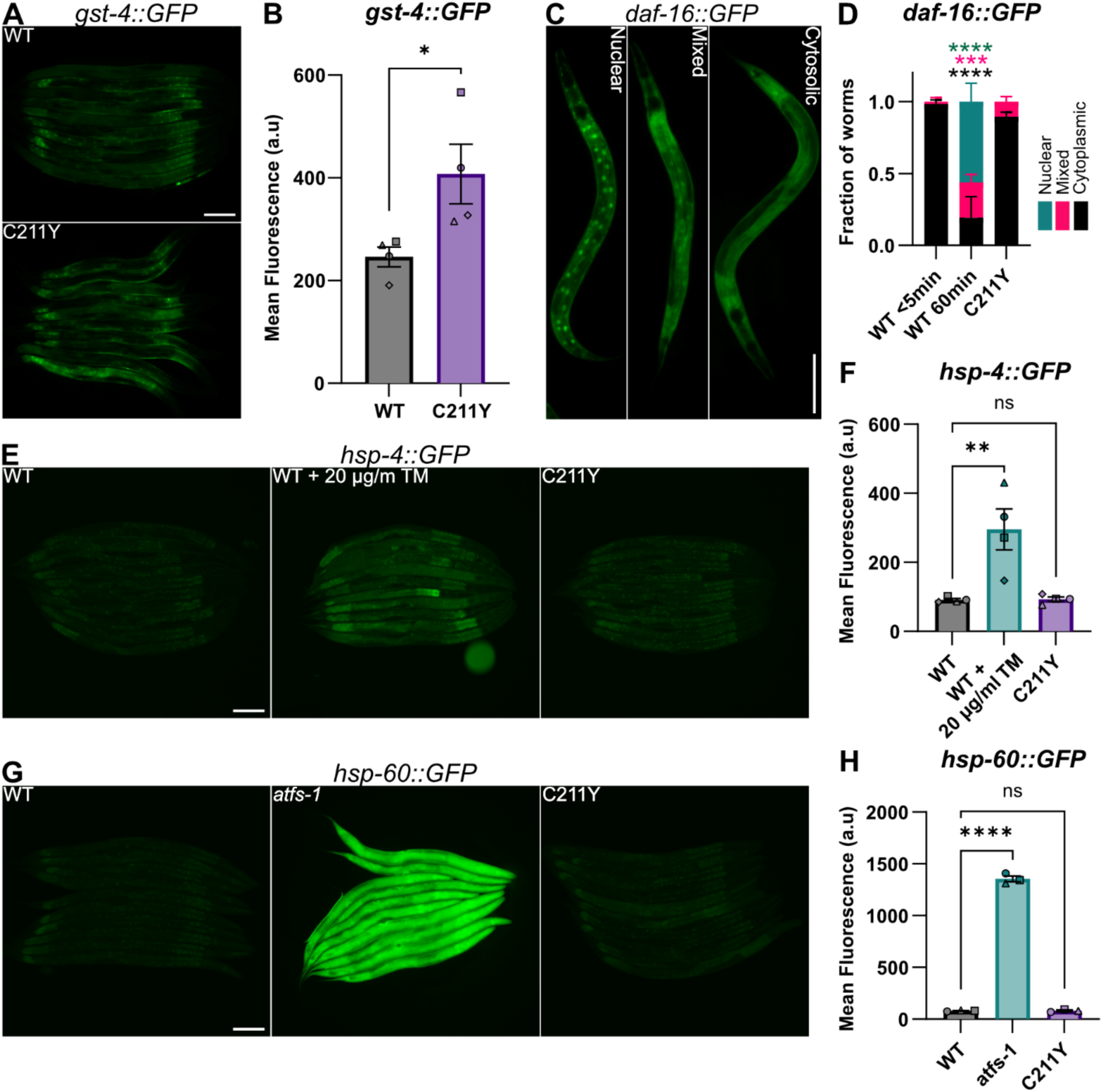
Expression of four stress reporters in the *pcyt-1(C211Y)* mutant. **A, C, E)** representative fluorescence images of worms expressing fluorescent ER-UPR (A), mt-UPR (C), and oxidative stress (E) reporters. **B, D,F)** Quantification of mean fluorescence in whole worms in A,C, and E respectively. **G)** Representative images of the three categories assigned to the *daf-16::GFP* reporter strain: Nuclear, Mixed and Cytoplasmic. **H)** Quantification of *daf-16::GFP* localization in G. Statistical analyses were done using Ordinary one-way ANOVA, with Dunnett’s multiple comparisons test for post-hoc analysis, n = 3 biological replicates (19–25 worms per replicate). * == p<0,5; ** == p<0,05; *** == p<0,005; **** == p<0,0005.

### P154A is mislocalized and has reduced protein levels at 25°C

Addition of a mNeonGreen tag at the C-terminal end of the endogenously-encoded PCYT-1 protein is well tolerated and allows detection of the protein in living worms at 20°C and 25°C, where it is enriched in the perinuclear regions most notably in intestinal cells but also in nuclei of other cells, for example epithelial cells where perinuclear localization can be seen in the hypodermal cells at the lateral sides of the worm, seen near the top and bottom focal planes in the z-stacks (**Fig. 5A; Suppl Movies 1-4**). A rather strong signal can be seen in the intestine for the WT worms, likely caused by gut granules, however in P154A this signal is weaker and the foci are less abundant than in WT (**Fig 5A**). In the P154A allele, we see one main differences at 25°C besides the difference in intestinal fluorescence: the signal from head nuclei is nearly invisible, suggesting reduced expression compared to the wild-type control (**Fig. 5A**). When whole-worm fluorescence of the mNeonGreen-tagged PCYT-1 is measured 24h after spotting L1s, we find that the fluorescence of the wild-type protein is higher than in P154A at 25°C, and that there is no difference in mean fluorescence between P154A at the two temperatures (**Fig. 5B-C**). Western blots quantifying protein abundance of endogenously-encoded 3xFLAG-tagged PCYT-1 48h after spotting L1s shows that control worms have a similar amount of protein as P154A mutants at 20°C, and that the amount of protein is lower in the P154A mutant when grown at 25°C (**Fig. 5D-E)**. In a separate quantification, we determined the ratio of nucleus-to-cytoplasm fluorescence in the first intestinal ring and found that this ratio is the same for the wild-type variant at 20°C and 25°C but reduced at 25°C for the P154A variant (**Fig. 5F-G**). The same ratio was also measured in the last intestinal ring where only P154A at 25°C had a reduced ratio. (**Fig. 5H-I).** Thus, both measurements are consistent with the temperature-sensitive nature of this allele since the reaction of this allele to 25°C differs from that of the wild-type protein. The decrease in non-nuclear fluorescence in P154A may suggest a decrease in gut granule formation or localization of the protein outside of the nucleus.

**Fig. 5.**
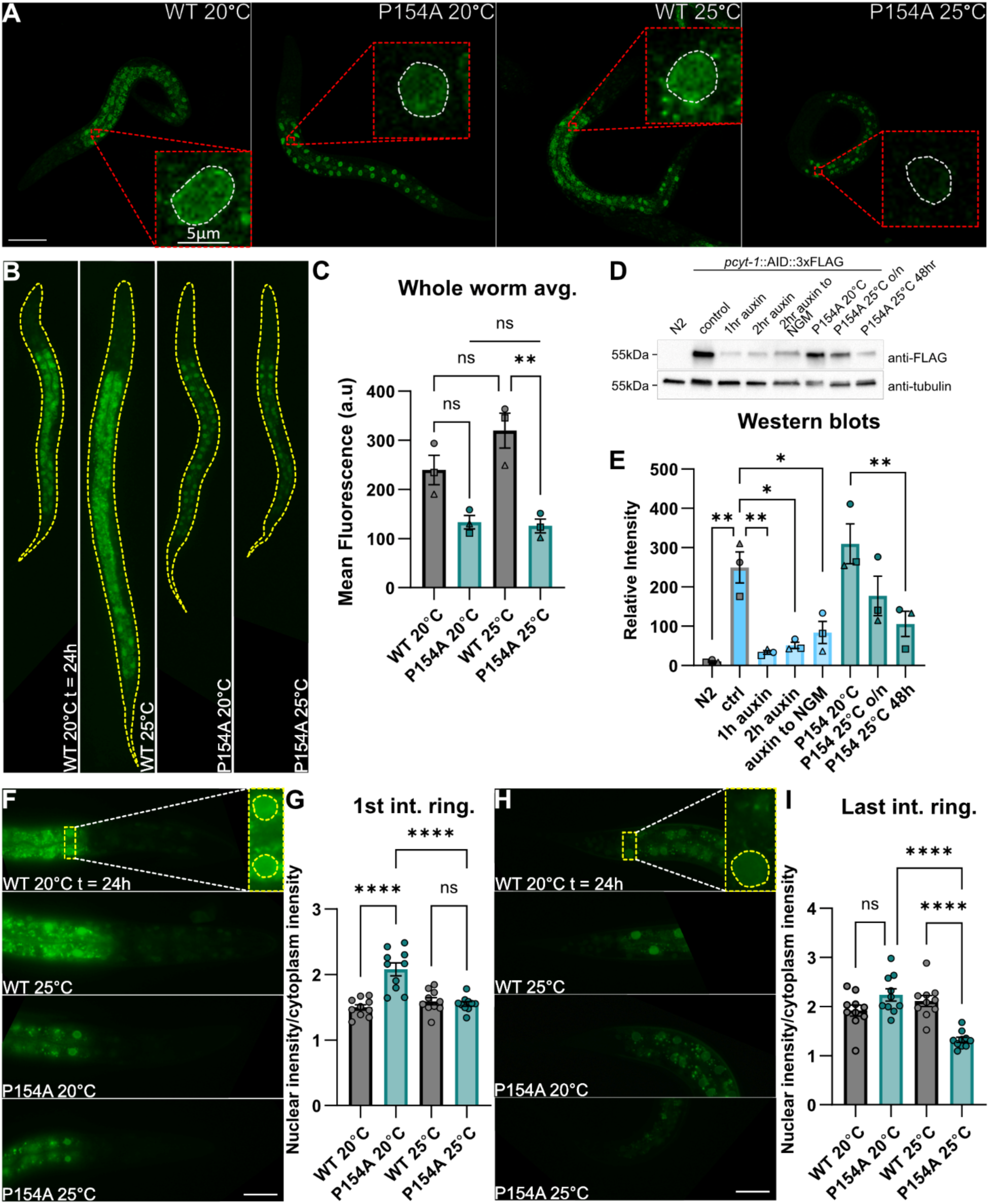
PCYT-1(P154A)::mNeonGreen localization and western blot quantification of PCYT-1 (P154A)::AID::3XFLAG and PCYT-1::AID::3XFLAG. **A)** Z-projections of z-stack images of *pcyt-1::mNeongreen* taken using a confocal microscope 24h after spotting L1s. Scalebar = 50um. Dashed red square shows an enlarged hypodermal cell nucleus in a single slice in the z-stack; white dashed lines outline the nucleus. **B)** Brightfield fluorescence images of *pcyt-1::mNeongreen* worms 24h after spotting L1s, used to quantify mean fluorescence of the whole worm. Dashed yellow line outlines the area measured. **C)** Quantification of worms in B. Ordinary ANOVA with Tukey’s multiple comparisons test, n = 3 biological replicates with ≥15 worms per replicate **D)** Representative western blot of proteins from worms with 3xFLAG and AID tagged PCYT-1. Auxin activates the degradations of AID tagged proteins. **E)** Quantification of western blots; the statistics was done using ordinary one-way ANOVA and Dunnett’s multiple comparisons test, n = 3 independent experiments. **F)** Representative images of head and first part of intestine that were used to measure the mean fluorescence of nuclei in the 1st intestinal ring and surrounding area. Dashed yellow line outlines the area measured. **G)** The ratio of the mean fluorescence in the nuclei and the surrounding cytoplasm shown in F. **H)** Representative images of tail and last part of intestine that were used to measure the mean fluorescence of nuclei in the last intestinal ring and surrounding area. Dashed yellow line outlines the area measured. **I)** The ratio of the mean fluorescence in the nuclei and the surrounding cytoplasm shown in H. For the quantification of nuclear to cytoplasmic ratio the statistics was done using ordinary one-way ANOVA and Dunnett’s multiple comparisons test, n = 10 worms

### Lipidomics confirms the *pcyt-1* allele hierarchy

Next, we analyzed the fatty acid composition of the most abundant phospholipids in *C. elegans,* namely the PCs and PEs, in L4 worms cultivated at 20°C or 25°C. The lipidomics analysis for worms grown at 20°C shows that the PCs of *pcyt-1* mutants have elevated levels of the LCPUFAs C20:3, C20:4 and C20:5 fatty acids at the expense of several shorter, more saturated FAs (**Fig. 6A**). Conversely, in PEs, the *pcyt-1* mutants show increased levels of C17:0 (likely the dietary C17iso) and decrease in C16:0 and C18:0 (**Fig. 6B**). The FA composition of the mutants was used in a principal component analysis (PCA), revealing a hierarchy of *pcyt-1* alleles at 20°C: N2<A97T<P154A<C211Y. Similar lipidomics findings were obtained at 25°C with the exception that the P154A allele is now as severe as C211Y (**Suppl. Fig. S1A-B**, **Fig. 6C**), confirming its temperature-sensitive nature. Interestingly, the PC/PE ratio is not affected in mutants grown at 20°C but is clearly reduced in the P154A and C211Y mutants at 25°C (**Fig. 6D**), suggesting that both mutants may be more severe at that temperature, which is also consistent with their worsened phenotypes (e.g., increased sterility) at that temperature. The FA composition data from two independent sets of experiments (each containing independent biological quadruplicates) at 20°C and 25°C is summarized in a heat map (**Fig. 6E**); note the general increase in LCPUFAs in the *pcyt-1* mutants, especially in C211Y at 20°C and in both C211Y and P154A at 25°C. The fatty acid composition changes are reflected in a direct comparison of the whole PCs and PEs between N2 and C211Y at 20°C: the most increased phospholipids in the mutant are PC19:1/20:5, PC18:1/20:5 and PE17:0/17:0, while the most decreased are PE 18:0/18:1, PE 16:0/18:1 and PC 18:1/PC18:1 (**Suppl. Fig S1C**). A general conclusion from this analysis is that the abundance of LCPUFAs are increased in the PCs and PEs at the expense of shorter FAs, with some exceptions: C17:0 (likely C17iso) is increased in mutants in both PCs and PEs, C20:5 is actually decreased in the PEs, and C19:1 (likely the cyclopropane C19delta) is clearly increased only in the C211Y mutant.

**Fig. 6.**
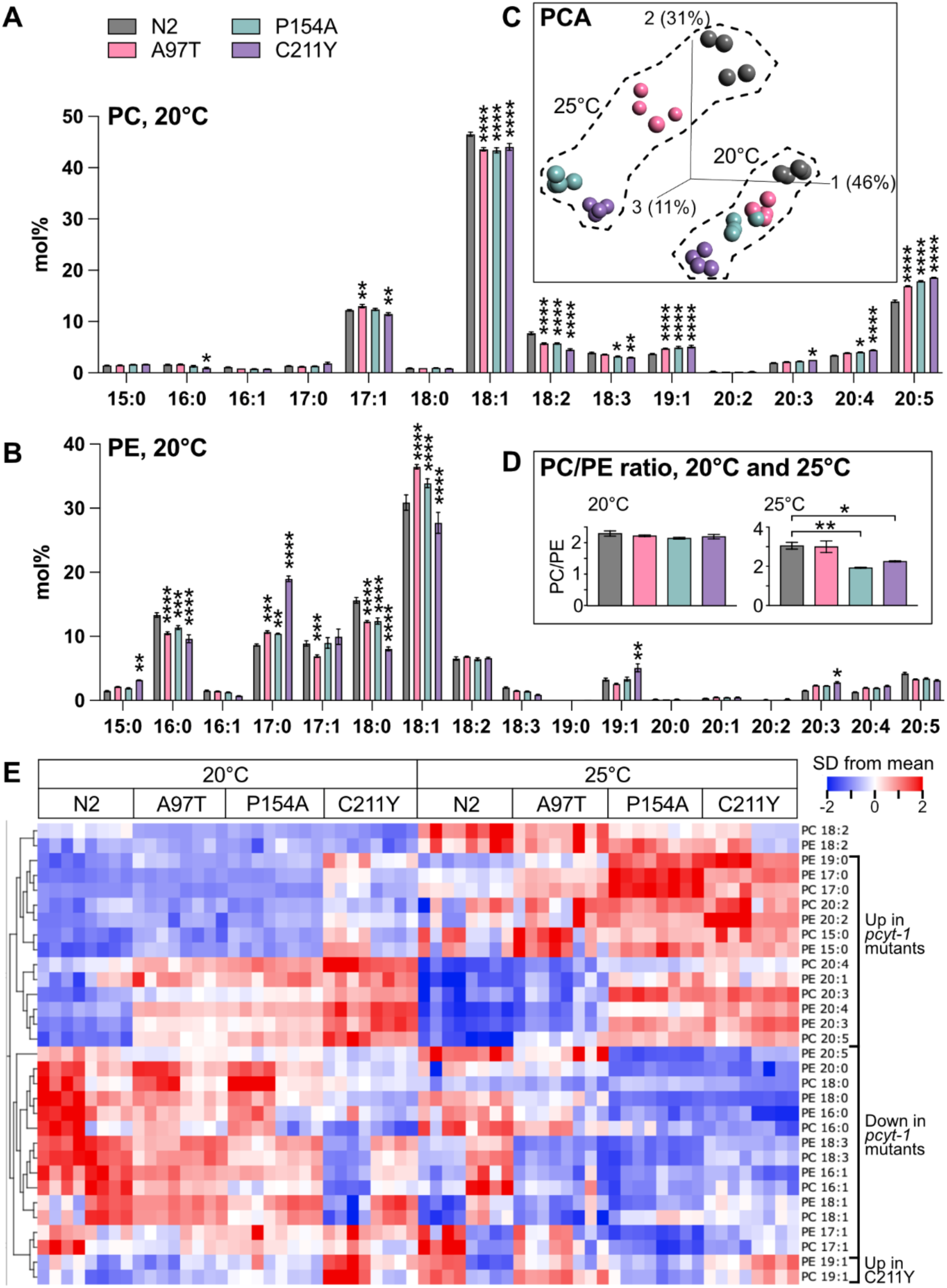
Lipidomics reveals a hierarchy of *pcyt-1* allele severity at 20°C and 25°C. **(A)** and **(B)** Show the fatty acid composition of phosphatidylcholines and phosphatidylethanolamines of L4 worms grown at 20°C and of the indicated genotypes; mean ± sem from four replicates is shown. **(C)** Shows a principal component analysis of the samples at 20°C and 25°C. **(D)** PC/PE ratios at 20°C and 25°C. **(E)** Heat map showing hierarchically clustered fatty acids; two independent sets of biological quadruplicates are shown for each condition. Several fatty acids are increased/decreased in the *pcyt-1* mutants (indicated at right) and suggest an allelic series at 20°C (N2<A97T<P154A<C211Y) and that P154A is a temperature-sensitive allele, resulting in this series at 25°C (N2<A97T<C211Y<P154A). For this analysis, the mol% fatty acid composition of phosphatidylcholines and phosphatidylethanolamines was determined using UPLC-MS/MS. For heat map, the mean mol% of each fatty acid among the samples was determined and set to 0 (zero) and the variance adjusted to 1, thus giving equal weight to all fatty acids. Note that the 15:0, 17:0 and 19:0 species likely consist mostly of mmBCFAs while the 17:1 and 19:1 species are likely mostly the dietary cyclopropanes 17:0 delta and 19:0 delta, respectively.

### Auxin-inducible degradation of PCYT-1

To complement the temperature-sensitive P154A allele, we also created an inducible loss-of-function *pcyt-1* allele using the auxin-inducible degradation (AID) system (Sharma *et al*. 2024). The efficacy of the AID system was evaluated using Western blots: addition of auxin to worms that globally express the *Arabidopsis* ubiquitinase TIR1 caused near complete degradation of PCYT-1::AID::3XFLAG within 1 hour, with minimal recovery after 2 hours of auxin treatment followed by an overnight cultivation on auxin-free plates (**Fig. 5E-F**). Lipidomics analysis of PCs and PEs reveals that auxin treatment by itself causes some changes in the FA composition of phospholipids, but without affecting the PC/PE ratio (**Fig. 7A-D**). Addition of auxin to PCYT-1::AID::3XFLAG causes a significant decrease in the PC/PE ratio, accompanied by many of the same changes previously observed with C211Y or P154A at 25°C (**Fig. 7A-D**), a general conclusion being again that the abundance of LCPUFAs are increased in the PCs and PEs at the expense of shorter FAs.

**Fig. 7.**
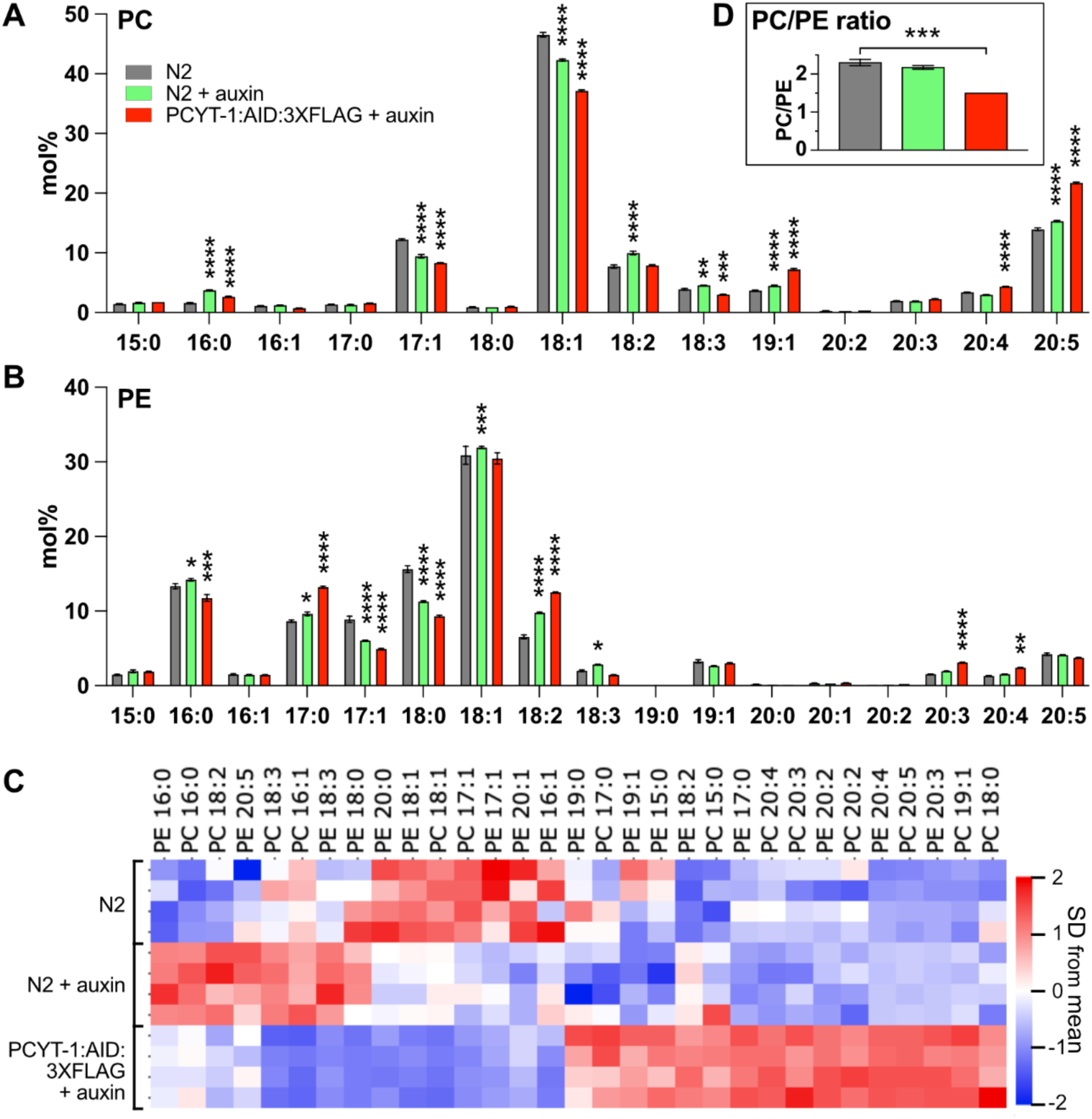
Lipidomics consequences of auxin-induced degradation of PCYT-1::AID::3XFLAG. **(A)** and **(B)** Show the fatty acid composition of phosphatidylcholines and phosphatidylethanolamines of L4 worms of the indicated genotypes and growth at 20°C with/without auxin for 6 hours. **(C)** The heat map shows the fatty acids clustered hierarchically based on similarity of mol% across treatments. For this analysis, the mol% fatty acid composition of phosphatidylcholines and phosphatidylethanolamines was determined using UPLC-MS/MS. For each fatty acid, the mean mol% among the samples was determined and set to 0 (zero) and the variance adjusted to 1, thus giving equal weight to all fatty acids. Note that the 15:0, 17:0 and 19:0 species likely consist mostly of mmBCFAs while the 17:1 and 19:1 species are likely mostly the dietary cyclopropanes 17:0 delta and 19:0 delta, respectively.

### Disrupting PC synthesis in sexually mature worms disrupts oogenesis

Others have described the sterility phenotype of *pcyt-1* mutants and suggested that it is caused by increased ferroptosis (Zhu *et al*. 2023), which is consistent with the increased peroxidation-prone LCPUFA levels that we observed. Here, we used the AID system to determine if the sterility defect in PCYT-1-deficient worms is strictly developmental or, alternatively, can be induced later in life. We found that inducing PCYT-1 degradation with auxin in L4 worms expressing PCYT-1::AID::3XFLAG and TIR1 leads to severe disruption of oogenesis within 24 hours, evidenced by a reduced number of maturing oocytes (made visible using a PIE-1::GFP reporter; (Audhya *et al*. 2005)) and aged or absent embryos in the uterus (**Fig. 8A-C**). Additionally, treating L1 worms expressing PCYT-1::AID::3XFLAG and TIR1 with auxin leads to immediate developmental arrest such that these worms are still L1s 48 hours later (**Fig. 8D**). We conclude that PCYT-1 is constantly required to sustain oogenesis and development, suggesting a steady demand for new synthesis of phosphatidylcholine in both processes.

**Fig. 8.**
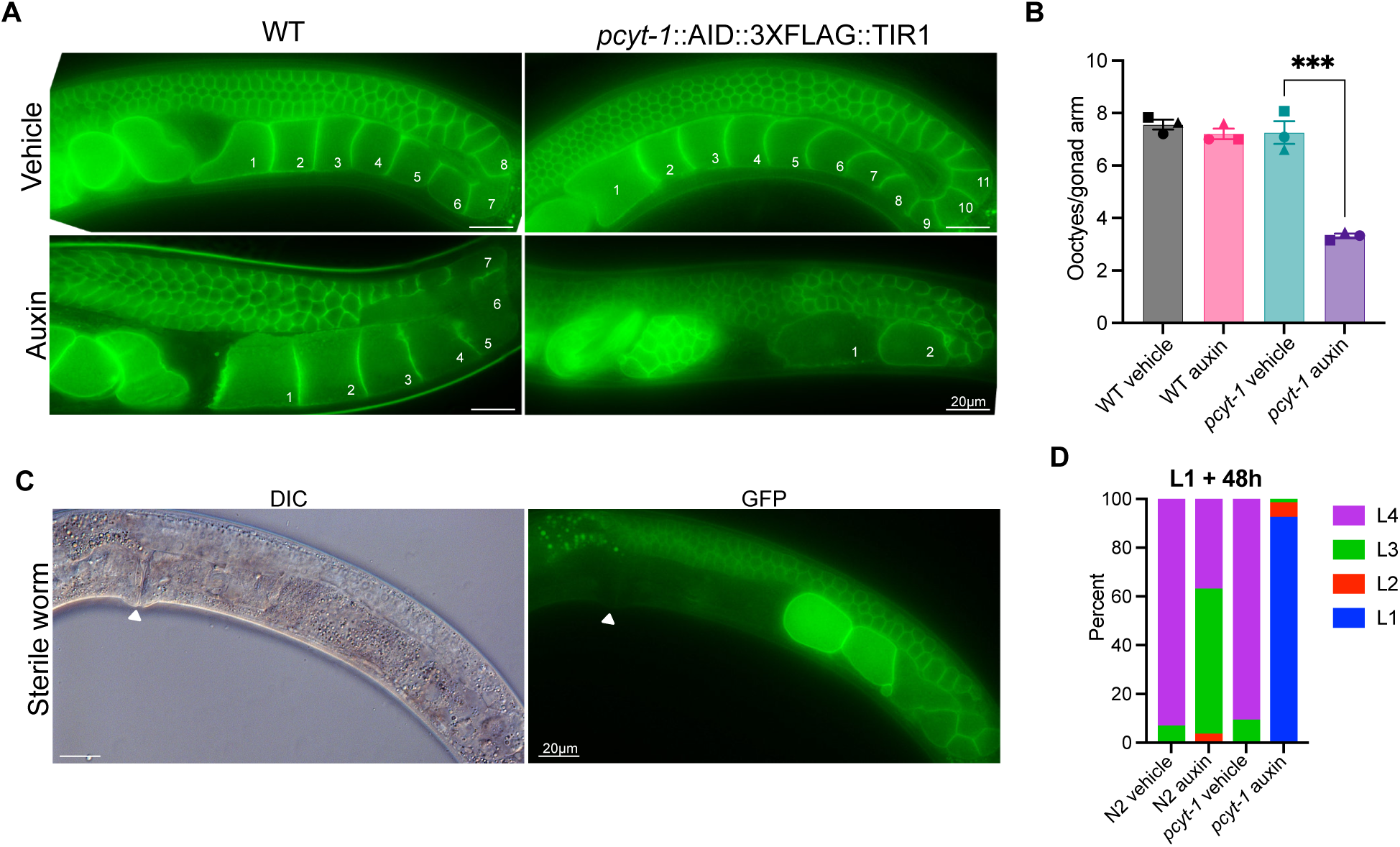
Auxin-induced PCYT-1::AID::GFP degradation causes germline and growth defects. **(A)** PIE-1::GFP shows that the organization of the germline and the number of maturing oocytes (numbered) are both impaired when *pcyt-1::AID::3XFLAG* L4 worms that also carry *eft-3P::TIR1* are treated with auxin. **(B)** Number of maturing oocytes per gonad arm in L4 worms cultivated overnight on control or auxin-containing plates. WT refers to N2 worms while *pcyt-1* refers to *pcyt-1::AID::3XFLAG* worms that also carry *eft-3P::TIR1* that supports expression of TIR1 in the soma. **(C)** Example of a sterile worm, with position of the vulva indicated in the DIC image and in the corresponding image showing the PIE-1::GFP signal. Note that the uterus is void and that the two oocytes in the proximal gonad arm express abnormally high levels of the PIE-1::GFP reporter. **(D)** Treating *pcyt-1::AID::3XFLAG* L1 larvae that also carry *eft-3P::TIR1* causes L1 arrest, as observed by scoring the developmental stages 48 hours later. Treatment of N2 worms with auxin alone also causes a slight developmental delay.

## DISCUSSION

The present study establishes a hierarchy among four point mutants in *C. elegans* PCYT-1: V146M (lethal)>C211Y (strong effects at 20°C, stronger at 25°C)>P154A (effects only at 25°C)>A97T (very mild). The C211Y variant causes several defects: slow growth, increased levels of LCPUFAs in phospholipids, upregulation of the oxidative stress reporter *gst-4::GFP*, oogenesis defects accompanied by sterility, suppression of *paqr-2* mutant phenotypes and, surprisingly, extended lifespan. Many of these defects were also found in the P154A mutant at its non-permissive temperature (i.e. 25°C). The C211Y sterility defects can be completely suppressed by providing choline, CDP-choline or lecithin (rich in phosphatidylcholine) in the diet, suggesting that it is a partial loss-of-function allele rather than some sort of neomorph. As mentioned in the introduction, three of the four alleles studied cause different effects in humans: A93T cause retinal dystrophies (Testa *et al*. 2017), V142M cause lipodystrophy and fatty liver disease (Payne *et al*. 2014) and P150A cause spondylometaphyseal dysplasia (a form of dwarfism) and cone-rod dystrophy. (Hoover-Fong *et al*. 2014; Wong 2014; Yamamoto *et al*. 2014). No C211Y orthologous variant has been described in human.

Given that PCYT-1 is responsible for the rate-limiting step in PC synthesis, it seems likely that the primary defect in the mutants relates to phospholipid composition, with the other defects being secondary. In particular, it is possible that a reduced rate of PC synthesis leads to a reduction in the PC turnover rate, allowing the Lands cycle more time to replace the FA at the sn-2 position with LCPUFAs, explaining their increased abundance in the *pcyt-1* mutants. The germline loss in *pcyt-1* mutants may be a secondary consequence of this caused by catastrophic ferroptosis to which the membrane network within the gonad becomes more sensitive as it accumulates LCPUFAs that can propagate lipid peroxidation (Perez *et al*. 2020).

Worms homozygous for the C211Y or P154A (at 25°C) variant have extended lifespan. This could be a direct result of the sterility since ablation of the germline extends lifespan in *C. elegans* (Hsin and Kenyon 1999; Lin *et al*. 2001; Arantes-Oliveira *et al*. 2002).

Alternatively, it is possible that reduced PC synthesis leads to the activation of protective pathways. In particular, lowering PC levels leads to ER-UPR activation and upregulation of lysosome-dependent proteostasis that contributes lifespan extension (Tong *et al*. 2026). However, an unexpected observation in our study is that the C211Y variant, which is severe enough to impact growth rate, brood size and lifespan at 20°C, does not trigger a strong mitochondrial, ER or metabolic stress response as assessed using commonly used GFP reporters. This suggests that reduced rate of PC synthesis is not per se causing stress within the cells even though it does lead to measurable changes in the FA composition of phospholipids. Only the GST-4::GFP reporter of oxidative stress response was upregulated in the C211Y mutant, consistent with the observed increase in lipid peroxidation-prone LCPUFAs in phospholipids.

That the fluorescently tagged P154A allele has a weaker fluorescent signal in the gut can have several different causes. The mutant protein can have trouble localizing to the cytoplasm or other membrane structure outside the nucleus, or it has fewer gut granules. The tagged protein has a rather weak fluorescence meaning that autofluorescence of gut granules are likely to contribute significantly to the fluorescent signal in the intestines. The potential decrease in gut granules in the P154A mutant may indicate that a reduction in PC synthesis or a defective PCYT-1 protein plays a role in gut granule formation. Gut granules are lysosome-related organelles known to play a role in storing trace metals and for synthesizing signaling molecules, however their function and development is still largely unknown (Hermann *et al*. 2005). It would therefore be of interest to investigate whether or not gut granule formation is affected in *pcyt-1* mutants and the mechanism behind this.

Reduced PC levels can cause activation of *sbp-1 (Walker et al. 2011)*, a transcription factor that contributes to the upregulation of desaturases, and this may be a mechanism by which C211Y allele acts as a *paqr-2* suppressor. However, here we show that the PC/PE ratio is not altered in the C211Y mutant grown at 20°C. Our results now suggest a different or complementary hypothesis: reduced PC synthesis leads to reduced PC turnover hence gradual accumulation of LCPUFAs in phospholipids via the Lands cycle. Further studies, such as careful measurements of SBP-1 and FAT-6 levels, and of lipid turnover rates in the C211Y, could help to clarify the exact mechanism of *paqr-2* suppression.

Finally, we showed that the PCYT-1::AID::3XFLAG provides an excellent way to experimentally induce PCYT-1 degradation, i.e. by addition of auxin to the culture plates, and that this also causes sterility in adult worms, which shows that the sterility is not the result of some developmental defect earlier in the life cycle. PCYT-1::AID::3XFLAG and the P154A variant thus offer two different ways to control PCYT-1 activity in worms, namely by the addition of auxin or by shifting to 25°C, respectively. These new genetic tools should facilitate further studies on the roles of PC homeostasis.

## MATERIALS AND METHODS

### Worm strains

The following strains were used in this study: the *C. elegans* wild-type strain N2, *paqr-2(tm3410)*, *pcyt-1(et9)*, *atfs1(et15)*, SJ4005 (*zcIs4 [hsp-4::GFP])*, SJ4058 (*zcIs9 [hsp-60::GFP +lin-15(+)])*, TJ356 (*zIs356 [daf-16p::daf-16a/b::GFP +rol6(su1006)]*), CL2166 (*dvIs19 [(pAF15)gst-4p::GFP::NLS]*), OD95 (*ltIs38 [pie-1p::GFP::PH(PLC1delta1) + unc-119(+)]), CA1200 (ieSi57 [eft-3p::TIR1::mRuby::unc-54 3’UTR + Cbr-unc-119(+)] II),* and are available from the *C. elegans* Genetics Center (CGC; USA). The strains PHX9942 (*pcyt-1::mNeongreen*) and PHX10600 (*pcyt-1::AID::3xFLAG*) were created by Suny Biotech Co using CRISPR/Cas9.

Worms were cultivated on NGM with *E. coli* OP50 as food source at 20°C unless otherwise stated. OP50 was maintained on LB plates at 4°C and re-streaked every 6-8 weeks. Single colonies were picked and cultivated in LB medium at 37°C overnight before being used to seed NGM plates.

### CRIPSR/Cas9 genome editing

Creation of *C. elegans* mutants with SNPs at loci homologous to missense mutations in human disorders was done using CRISPR/Cas9 gene editing as previously described (Dokshin *et al*. 2018). Alt-R HDR Donor Oligo ssDNA, Alt-R™ CRISPR-Cas9 crRNA, Alt-R™ CRISPR-Cas9 tracrRNA, and Alt-R™ S.p. Cas9 Nuclease V3 protein were ordered from IDT (Integrated DNA Technologies, Inc; Coralville, IA, USA). ssDNA and crRNA were designed using IDTs Alt-R™ HDR Design Tool. Creating the *pcyt-1* missense mutants A97T, V142M, and P154A was done using the crRNA sequences 5’-GUUGAUUCGCAUGCCCAUGAGUUUUAGAGCUAUGCU-3’, 5’-ACGAUGGUGUUCGGCACUGUGUUUUAGAGCUAUGCU-3’, 5’-AAGUCUACAGAGAGGCUCCAGUUUUAGAGCUAUGCU-3’ and the ssDNA donor sequences 5’-TGCGAATTTATGCTGATGGAATTTATGATCTGTTCCACCATGGGCATACGAACCAACTTCGCCAAGTGAAGAAAATGTTCCCAAATGTCT-3’, 5’-AGTCACTTCAGAAGAAGAACGTTACGATGGTGTTCGGCACTGCCGTTACATGGATGAAGTCTACAGAGAGGCTCCATGGTTTTGTACTGTCG-3’, and 5’-TCGGCACTGTCGGTACGTTGATGAAGTCTACAGAGAGGCAGCATGGTTTTGTACTGTCGAGTTTTTGAAAAACCTTAAAGTTG-3’ respectively. The injection mixes were prepared using 10 μg/μl of the Cas9 enzyme (IDT), 0.4 μg/μl tracrRNA (IDT), 0.4 μg/μl crRNA (IDT), 1 μg/μl of ssDNA (IDT), and 40 ng/μl of *Pmyo-2(GFP)* plasmid. The mixture was microinjected into the gonad of day 1 adult worms and the F1 generation was screened for animals expressing the reporter plasmid. Genotypes were tested by PCR and successfully edited genes were confirmed by Sanger sequencing (Eurofins). The missense mutants were created by injecting N2 worms. The P154A missense mutation was also introduced into the strains carrying *pcyt-1::mNeonGreen* and *pcyt-1::AID::3xFLAG*.

### Phenotypic assays

For all phenotypic assays worms were synchronized by egg laying unless otherwise stated. Fertile adult worms were placed on the test plate to lay eggs for ≥2h, until ≥30 eggs were laid, the adult worms were then removed from the plate. For the growth assay, worms were imaged using a Zeiss Axioscope using a 10x objective after 72h at 20 and 25°C, or 144h at 15°C, and the lengths were measured using ImageJ (Schneider *et al*. 2012). Brood size was measured by placing L4 worms on the test plate on day 0. On every subsequent day for 8 days the parent worms were transferred to a new plate and the number of total hatched progenies was counted for each strain. Statistical analyses were done using a Two-way mixed-effects ANOVA (REML) using LS means (GraphPad Prism), with biological replicate treated as a random effect. Tukey’s multiple comparisons test was used for post-hoc analysis, comparing each strain to N2.

For the sterility assays, sterility was scored as absence of eggs in the uterus, and the fraction of sterile worms was counted after 96h at 20 or 25°C, or 168h at 15°C. Choline chloride (C7527 Sigma-Aldrich), CDP-choline (Cytidine 5′-diphosphocholine sodium salt hydrate, 30290 Sigma-Aldrich) and soy lecithin (L-α-lecithin, 429415 Sigma-Aldrich) were sterilized and added to the agar before pouring the test plates. Statistical analyses were done using Ordinary one-way ANOVA, with Dunett’s multiple comparisons test for post-hoc analysis comparing each strain to N2, or untreated control.

The lifespan assay was performed by picking L4s from a synchronized plate on day zero, twelve L4s per plate for ten plates per strain. Plates were examined for dead worms every day until all worms were dead. Worms were transferred every few days to avoid confusion with progeny. Statistical analyses were done using the log-rank (Mantel-Cox) test comparing each mutant to N2 individually for each experiment.

### Stress response assays

Worms were imaged using a Zeiss Axioscope with a 10x objective. For each stress reporter, n ≥ 3 replicates with ≥20 worms per replicate. For worm strains carrying *hsp-4::GFP*, *hsp-60::GFP*, and *gst-4::GFP*, worms were imaged as day 1 adults and the mean fluorescence intensity of whole worms were measured using ImageJ (Schneider *et al*. 2012). The statistical analyses were done using a Two-way mixed-effects ANOVA (REML) using LS means (GraphPad Prism), with biological replicate treated as a random effect. Tukey’s multiple comparisons test was used for post-hoc analysis. Worm strains carrying *daf-16::GFP* were also imaged as day 1 adults and the fraction of worms with cytoplasmic, mixed and nuclear localization was quantified. Statistical significance was tested using a Two-way ANOVA and Tukey’s multiple comparisons test for post-hoc analysis comparing within each outcome category.

### Protein localization

Worms carrying mNeonGreen tagged PCYT-1 were synchronized by bleaching and spotted as L1s on the test plate. The animals were imaged after 24h using a Zeiss Axioscope using a 20x objective and 63x oil immersion objective, or using a Zeiss LSM700inv laser scanning confocal microscope with a 40X water immersion objective. mNeonGreen was excited with a 488 nm laser and images were collected at an emission of 515 nm. Image analysis was done in ImageJ (Schneider *et al*. 2012). Mean fluorescence intensities were measured on whole worms, and the ratio of mean fluorescence intensity in the nucleus versus background was measured by selecting a rectangular area covering two nuclei of the first intestinal ring and the width of the intestine, and a rectangle covering the posterior-most nucleus in the last intestinal ring and the width of the intestine. Statistical testing was done using a Two-way mixed-effects ANOVA (REML) using LS means (GraphPad Prism), with biological replicate treated as a random effect. Tukey’s multiple comparisons test was used for post-hoc analysis.

### Auxin treatment

Natural auxin indole-3-acetic acid (IAA) was added to molten NGM agar prior to pouring plates for a final concentration of 4 mM auxin. As auxin inhibits bacterial growth, fresh OP50 culture was concentrated 10X before being seeded onto cooled plates. For PCYT degradation experiments, synchronized worms with an endogenous tagged PCYT-1::AID::3XFLAG expressing somatic *Arabidopsis* TIR1 were transferred to auxin plates as L1s or L4s and incubated for 48 h or 24 h respectively. Worms were mounted on slides and photographed. The developmental stage of worms was scored. Statistical analyses were done using Ordinary one-way ANOVA, with Tukey’s multiple comparisons test for post-hoc analysis.

### Western blots

Synchronized worms with an endogenous tagged PCYT-1::AID::3XFLAG and expressing somatic TIR1 were incubated on NGM plates for 46 h at 20°C before being transferred to plates containing ethanol (control) or auxin for 1 or 2 hours. Worms in the “auxin to NGM” sample were incubated for 30 h on NGM before being transferred to auxin plates for 2 h and then back to NGM plates for an addition 14 h. Additionally, synchronized worms with the *pyct-1(P154A)* mutation introduced into the PCYT-1::AID::3XFLAG strain by CRISPR-Cas9 were incubated at 20°C for 48 h, 25°C for 48 h, or 20°C for 32 h followed by 16 h at 25°C. Samples were washed 3 times with M9 and then lysed using lysis buffer containing 25 mM Tris (pH 7.5), 300 mM NaCl, 0.1% NP40, and 1X protease inhibitor on ice with a motorized pestle. Samples were centrifuged at 20000g for 15 min at 4°C and protein samples were quantified using BCA protein assay kit. 15 μg of protein was mixed with Laemmli sample loading buffer containing β-mercaptoethanol, boiled for 10 min, and loaded on a 4% to 20% gradient precast SDS gel. After electrophoresis, the proteins were transferred to nitrocellulose membranes using Trans-Blot Turbo Transfer Packs and a Trans-Blot Turbo apparatus and predefined mixed-MW program. Blots were blocked in 5% nonfat dry milk in PBST for 1 h at room temperature and incubated with primary antibodies (mouse monoclonal anti-FLAG (M2, Sigma Aldrich) 1:5000 dilution and mouse monoclonal anti-alpha-Tubulin (B512, Sigma Aldrich) 1:5000 dilution) for 1 h at room temperature. Blots were then washed with PBST and incubated with goat anti-mouse HRP 1:3000 dilution for 1 h at room temperature and washed again with PBST. Detection of the hybridized antibody was performed using ECL detection kit (Immobilon Western, Millipore) and the signal was visualized with a digital camera (VersaDoc, Bio-Rad).

### Lipidomics

Samples were composed of synchronized L4 larvae grown on NGM or auxin plates at 20°C or 25°C (one 9 cm diameter plate/sample). For each treatment/genotype, two completely independent sets of samples were prepared, each set containing four independently grown replicates. Auxin treated samples were grown on NGM plates until late L3/early L4 stage before being transferred to auxin plates 6 h before collection. Worms were washed 3 times in M9, pelleted, and stored at −80°C until further analysis.

For lipid extraction, the pellet was sonicated for 10 minutes in butanol:methanol [3:1] and then extracted according to published methods (Lofgren *et al*. 2016). The total lipid extracts were evaporated under a stream of nitrogen and reconstituted in butanol:methanol [3:1] containing a 1:200 dilution of the Equisplash internal standard mix (Avanti lipids). The extracts were injected (2µL) onto a Waters BEH C18 column (2.1×50mm; 1.7µm) and phospholipid species were separated using water:acetonitrile:isopropanol [50:30:20] as mobile phase A and water:acetonitrile:isopropanol [1:9:90] as mobile phase B. Both mobile phases contained 10mM of ammonium formate. The flowrate was 400 µL/min and the column was kept at 40°C. The phosphatidylcholines (PCs) and phosphatidylethanolamines (PEs) were monitored using negative electrospray ionization on a TQ Absolute (Waters). The PCs were detected as formate adduct while the PEs were detected as [M-H]^-^. In total, 127 MRM transitions were monitored, and relative quantification of the PC and PE species was made using the signal from the internal standard. The data was evaluated using the TargetLynx software (Waters).

Statistical significance in each experiment was tested using a Two-way ANOVA and Dunnett’s multiple comparisons test for post-hoc analysis comparing the abundance of each lipid between the genotypes. Qlucore Omics Explorer n.n (Qlucore AB) was used for PCA analysis, generation of a volcano plot and displaying the data on heat maps; the data were normalized for the purpose of the heat map visualization (mean = 0; variance = 1).

### Germline membrane morphology

L4 worms carrying the *ItIs38 [pie-1p::GFP::PH(PLCdelta1) + unc-119(+)]* transgene were transferred from NGM plates to ethanol (vehicle) or auxin containing plates and incubated overnight at 20°C. The worms were mounted on agarose pads and then imaged with a Zeiss Axioscope with a 63x oil immersion objective. Results were quantified by counting the number of developing oocytes visible in gonad arms (one gonad arm/worm was scored). Statistical analyses were done using Ordinary one-way ANOVA, with Tukey’s multiple comparisons test for post-hoc analysis

## ACKNOWLEDGEMENTS

This work was supported by the Swedish Research Council grants 2024-04012, Cancerfonden grant 25 4340 Pj 01 H, and Wilhelm och Martina Lundgrens Vetenskapsfond grant 2025-GU-5047. Some strains were provided by the CGC, which is funded by NIH Office of Research Infrastructure Programs (P40 OD010440). We acknowledge the Centre for Cellular Imaging at the University of Gothenburg and the National Microscopy Infrastructure, NMI (VR-RFI 2019-00217) for providing assistance in microscopy.

## Supplementary Figures

**Suppl. Fig. S1.**
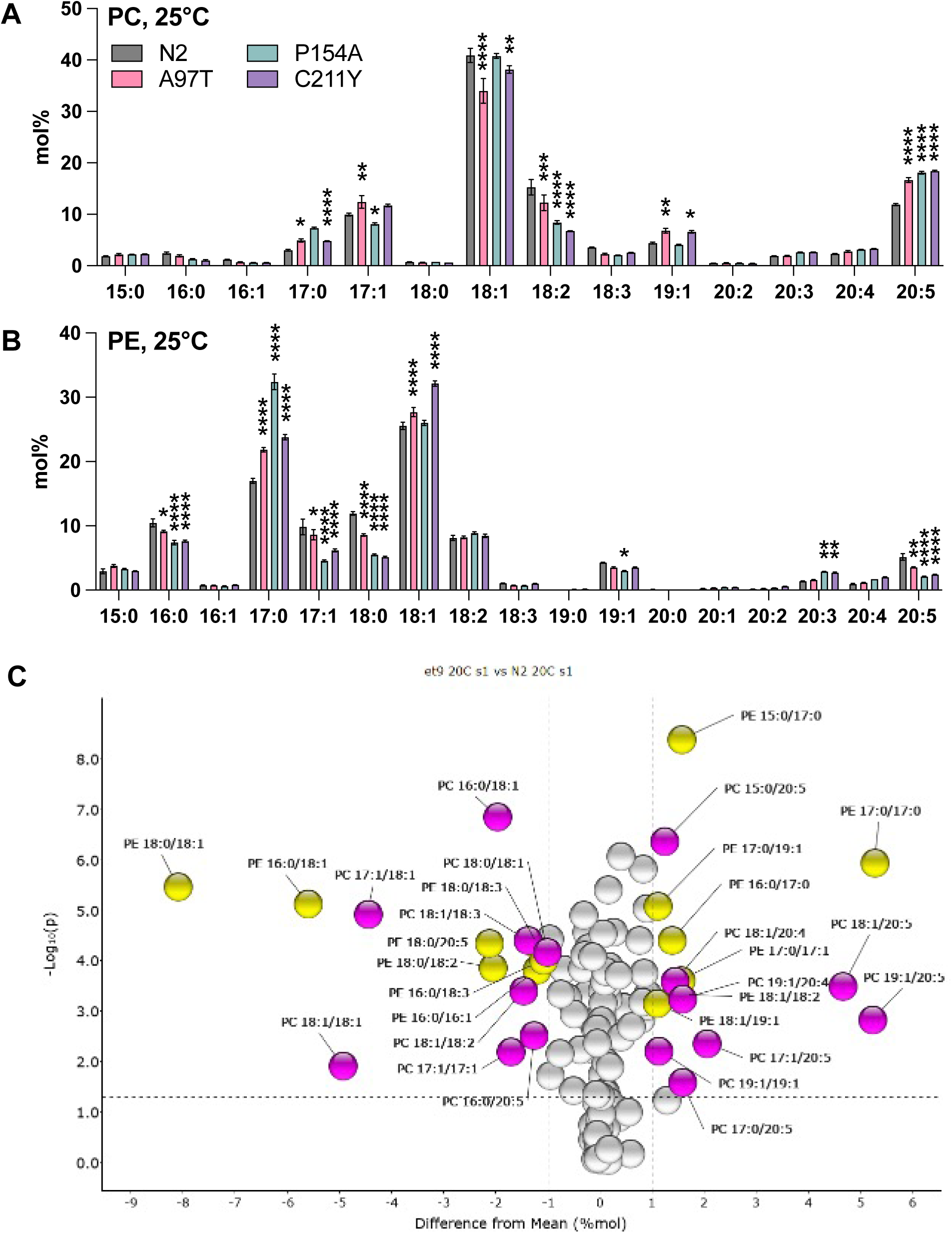
Lipidomics reveals a hierarchy of *pcyt-1* allele 25°C. **(A)** and **(B)** Show the fatty acid composition of phosphatidylcholines and phosphatidylethanolamines of L4 worms growth at 25°C and of the indicated genotypes; this data is included in the heat map from Fig. 6. For this analysis, the mol% fatty acid composition of phosphatidylcholines and phosphatidylethanolamines was determined using UPLC-MS/MS. For each fatty acid, the mean mol% among the samples was determined and set to 0 (zero) and the variance adjusted to 1, thus giving equal weight to all fatty acids. Note that the 15:0, 17:0 and 19:0 species likely consist mostly of mmBCFAs while the 17:1 and 19:1 species are likely mostly the dietary cyclopropanes 17:0 delta and 19:0 delta, respectively. **(C)** Volcano plot of PC and PE species in *pcyt-1(C211Y)* vs control N2 worms.

**Suppl Movies 1-4**

Z-stack of PCYT-1::AID::3XFLAG (WT for Movies 1 and 3; P154 variant for Movies 2 and 4) grown for 24 hours at either 20°C (Movies 1-2) or 25°C (Movies 3-4) post L1 stage.

